# Deep learning for early warning signals of regime shifts

**DOI:** 10.1101/2021.03.28.437429

**Authors:** Thomas M. Bury, R. I. Sujith, Induja Pavithran, Marten Scheffer, Timothy M. Lenton, Madhur Anand, Chris T. Bauch

## Abstract

Many natural systems exhibit regime shifts where slowly changing environmental conditions suddenly shift the system to a new and sometimes very different state. As the tipping point is approached, the dynamics of complex and varied systems all simplify down to a small number of possible ‘normal forms’ that determine how the new regime will look. Indicators such as increasing lag-1 autocorrelation and variance provide generic early warning signals (EWS) by detecting how dynamics slow down near the tipping point. But they do not indicate what type of new regime will emerge. Here we develop a deep learning algorithm that can detect EWS in systems it was not explicitly trained on, by exploiting information about normal forms and scaling behaviour of dynamics near tipping points that are common to many dynamical systems. The algorithm detects EWS in 268 empirical and model time series from ecology, thermoacoustics, climatology, and epidemiology with much greater sensitivity and specificity than generic EWS. It can also predict the normal form that will characterize the oncoming regime shift. Such approaches can help humans better manage regime shifts. The algorithm also illustrates how a universe of possible models can be mined to recognize naturally-occurring tipping points.

## Introduction

Many natural systems alternate between states of equilibrium and flux. This has stimulated research in fields ranging from evolutionary biology^1^ and statistical mechanics^2^ to dynamical systems theory^3^. Dynamical systems evolve over time in a state space described by a mathematical function^3^. Thus, dynamical systems are extremely diverse, ranging in spatial scale from the expanding universe^4^ to quantum systems^5^ and everything in-between^6–8^.

Different dynamical systems exhibit vastly different levels of complexity, and correspondingly diverse behaviour far from equilibrium states. Sometimes, a system that is close to equilibrium may experience slowly changing external conditions that move it toward a tipping point (‘bifurcation point’) where its qualitative behaviour changes. In these circumstance, dynamical systems theory predicts that even very high-dimensional systems will simplify to follow low-dimensional dynamics^9,10^. Moreover, all dynamical systems go through one of a limited number of possible bifurcation types, called ‘normal forms’ (Box 1)^3^. For instance, in a fold bifurcation, the system exhibits a critical transition–a sudden transition to a very different state. A transcritical bifurcation usually causes a smooth transition, though it may sometimes cause a critical transition^11^. Or, a Hopf bifurcation can lead the system into a state of oscillatory behaviour.

### Box 1. Dimension reduction close to a bifurcation.

As a high-dimensional dynamical system approaches a bifurcation, its dynamics simplify according to the centre manifold theorem^9^. That is, the dynamics converge to a lower-dimensional space, which exhibits dynamics topologically equivalent to those of the *normal form* of that bifurcation. Examples of a fold, (supercritical) Hopf and transcritical bifurcation are shown in Box 1 Figure a,b,c. Dynamics close to the bifurcation (grey box) are topologically equivalent to the normal forms

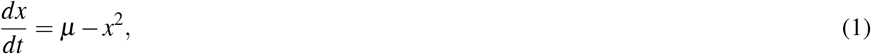

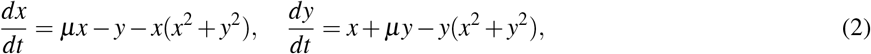

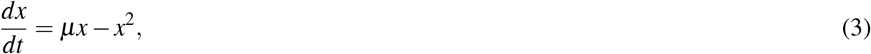

respectively, where *µ* is the bifurcation parameter. The bifurcation of each system occurs at *µ* = 0. These normal forms are contained within the set of 2-dimensional dynamical systems with third-order polynomial right-hand sides, motivating this as the framework for training the machine learning algorithm (see Methods).

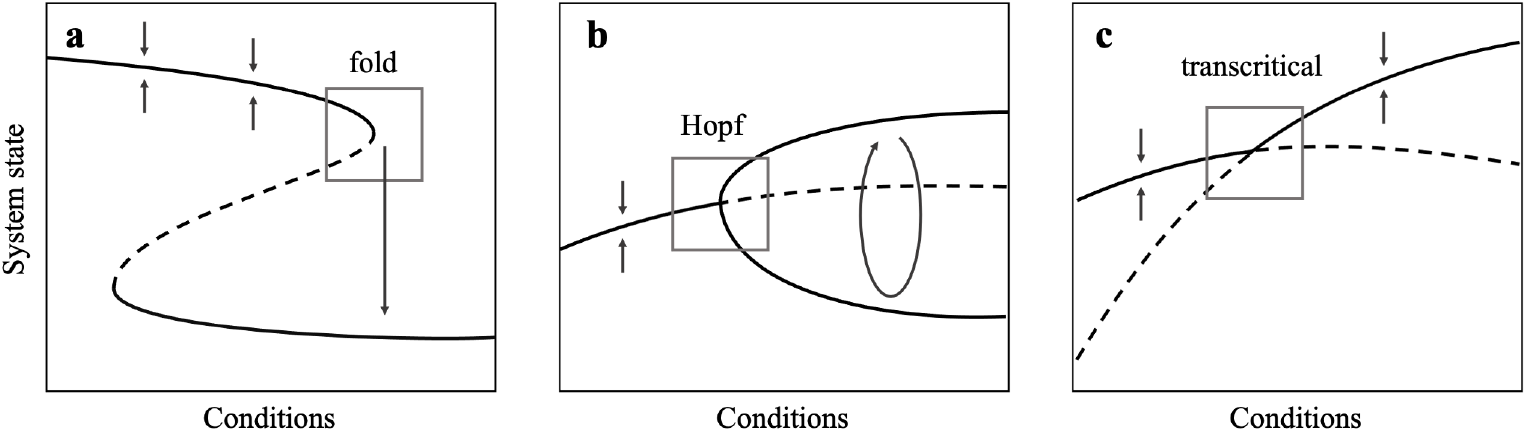

The normal form dictates how the system behaves near the bifurcation. However, other behaviours can emerge near the bifurcation that are common to all normal forms. For example, a system may undergo critical slowing down: system dynamics become progressively less resilient to perturbations as the transition approaches. This causes dynamics to become more variable and auto-correlated. As a result, statistical indicators such as rising variance and lag-1 autocorrelation (AC) of a time series often precede critical transitions in a variety of systems^12–14^. These generic early warning indicators have been found to precede critical transitions in systems including epileptic seizures, the Earth’s paleo-climate system, and lake manipulation experiments^15–17^.

Mathematically, critical slowing down occurs when the real part of the dominant eigenvalue (a measure of system resilience, Box 2) diminishes and eventually passes through zero at the bifurcation point. This happens for fold, Hopf and transcritical bifurcations and thus critical slowing down is manifested before all three bifurcations types^14^. Generic early warning indicators are intended to work across a range of different types of systems by detecting critical slowing down. But this strength is also their weakness, since these indicators do not tell us which type of bifurcation to expect^14^.

### Box 2. The significance of higher-order terms close to a bifurcation.

The local behaviour of a dynamical system is often well described by a linear approximation. However, for systems nearing a bifurcation, higher-order terms become significant. We illustrate this for a one-dimensional system *dx/dt* = *f* (*x*) with equilibrium *x*^∗^ i.e. *f* (*x*^∗^) = 0. The dynamics about equilibrium following a perturbation by *ε* Satisfy

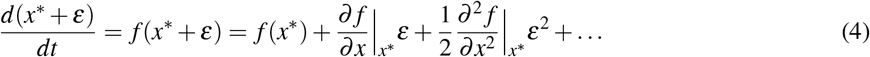

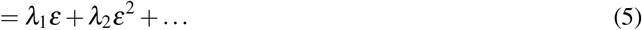

where *λ*_1_, *λ*_2_, … are coefficients of the Taylor expansion. The potential landscape centred on *x*^∗^ is given by

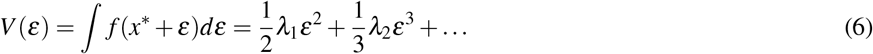

where we have dropped the arbitrary integration constant. Far from a bifurcation in a regime of small noise, displacement from equilibrium (*ε*) is small, and so the visited part of the potential landscape is well described by the first-order approximation (Figure a). As a local bifurcation is approached, *λ*_1_ → 0, which corresponds to critical slowing down, and a flattening of the first-order approximation to the potential landscape (Figure b). This allows noise to push the system further from equilibrium where higher-order terms become significant.

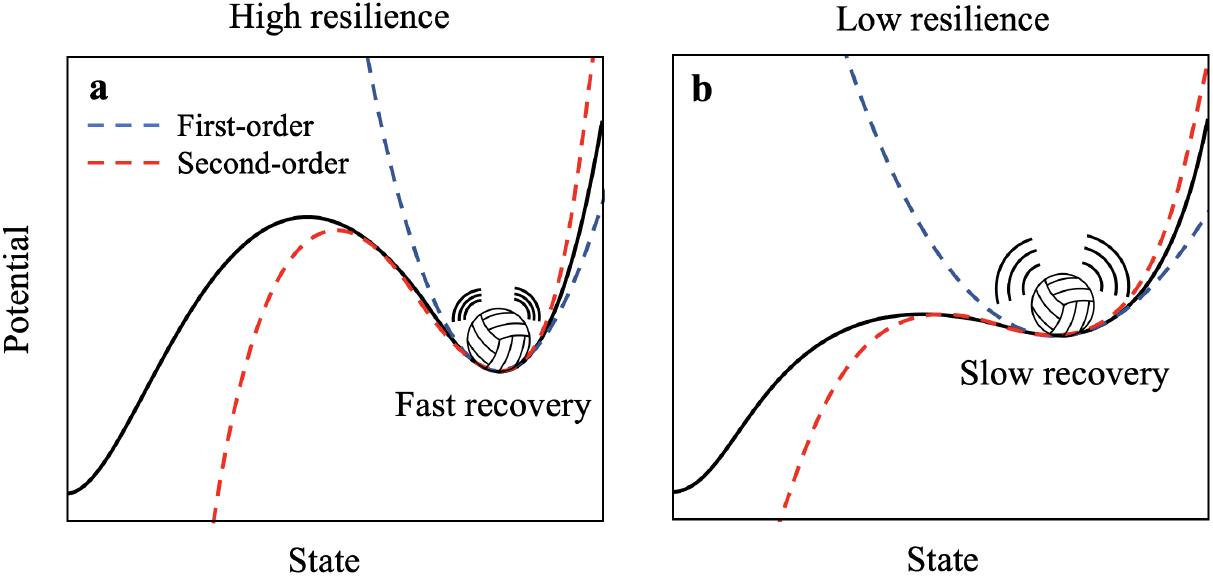

The dominant eigenvalue is derived from a first-order approximation to dynamics near the equilibrium. Higher-order approximations can distinguish between different types of bifurcations. But they are not often used to develop early warning indicators because (1) the first-order approximation dominates dynamics sufficiently close to the equilibrium, causing critical slowing down to generate the strongest signal, and (2) the first-order approximation is more tractable to mathematical analysis of stochastic systems than the higher-order approximations^18^. However, as a system gets closer to a bifurcation it can drift further from equilibrium due to critical slowing down. As a consequence, the higher-order terms become significant and may be large enough to provide clues about the type of transition that will occur. Statistical measures such as skew and kurtosis reflect the influence of these highest order terms, for instance^18–20^. Higher-order terms could be associated with features in time series data that are subtle but detectable, if we knew what to look for. Knowing whether a regime shift will be sudden or gradual–and what kind of new state will emerge beyond the tipping point–could be valuable in a range of applications.

### Deep learning and bifurcation theory

Generic early warning indicators such as variance and lag-1 AC use insights from dynamical systems theory to detect patterns that emerge before a regime shift (Box 2). Supervised deep learning (DL) algorithms can also detect patterns (features) in time series, and have achieved state-of-the-art in time series classification^21^ –the ability to classify time series based on characteristic features in the data. We hypothesised that DL algorithms can detect not only critical slowing down, but other subtle behaviour that occurs in time series prior to each type of bifurcation, such as the patterns generated by higher-order terms.

However, supervised DL algorithms require many thousands of time series to learn classifications–something we do not have for many empirical systems^22^. And they can only classify time series similar to the type of data they were trained on. Here, we propose that simplification of dynamical patterns near a bifurcation point provides a way to address the problem of limited empirical data, and moreover allows us to relax the restriction that DL algorithms can only classify time series from systems that they were trained on. Our first hypothesis (H1) is that if we train a DL algorithm on a sufficiently large training set generated from a sufficiently diverse library of possible dynamical systems, the relevant features of any empirical system approaching a regime shift will be represented somewhere in that library. Therefore, the trained algorithm will detect EWS in empirical systems that are not explicitly represented in the training set. Thus, even a relatively limited library might contain the right kinds of features that characterise higher-order terms in real-world time series. Our second hypothesis (H2) is that the DL algorithm will detect early warning signals with greater sensitivity and specificity (fewer false positives) than generic early warning indicators. Our third hypothesis (H3) is that the DL algorithm will also predict what type of regime shift is coming, on account of being able to recognise patterns associated with higher-order terms. All three hypotheses are based on the simplification of complex dynamics close to a bifurcation point (Boxes 1, 2).

To test these hypotheses, we developed a DL algorithm to provide early warning signals for regime shifts in systems it was not trained upon. We used a CNN-LSTM architecture (convolutional neural network—long short-term memory network, see Methods). CNN-LSTM sandwiches two different types of neural network layers. The CNN layer reads in subsequences of the time series and extracts features that appear in those subsequences. The LSTM layer then reads in the output of the CNN and interprets those features. The LSTM layer loops back on itself to generate memory, enabling the layer to recognize the same feature appearing at different times in a long time series. As a result, this this approach excels at pattern recognition and sequence prediction^23,24^

We created a training set consisting of simulations from a randomly-generated library of mathematical models exhibiting local bifurcations (see Methods). Specifically, we generated three classes of simulations eventually going through a fold, Hopf or transcritical bifurcation, and a fourth neutral class that never goes through a bifurcation. Then, we trained the CNN-LSTM algorithm on the training set to classify any given time series into one of the four classes based on the pre-bifurcation portion of the simulation time series. The *f*_1_ score of the algorithm tested against a hold-out portion of the training set was 88.2% when training on time series of length 1500 datapoints, and 84.2% when training on time series of length 500 datapoints.

We evaluated the out-of-sample predictive performance of the algorithm, using data from study systems that were not included in the training set. We tested three model systems and three empirical systems. The model systems included a simple harvesting model consisting of a single equation that exhibits a fold bifurcation^25^ ; a system of two equations representing a consumer-resource (predator-prey) system exhibiting both Hopf and transcritical bifurcations^26^ ; and a system of five equations representing the coupled dynamics of infection transmission and vaccine opinion propagation, and exhibiting a non-smooth transcritical bifurcation^11^. The three empirical datasets consisted of data from paleo-climate transitions^15^ ; transitions to thermoacoustic instability in a horizontal Rijke tube which is a prototypical thermoacoustic system^27^ ; and sedimentary archives capturing episodes of anoxia in the eastern Mediterranean^28^ (see Methods). We selected these empirical datasets because in each case they have been previously argued to show critical slowing down based on lag-1 AC and/or variance trends, followed by observation of a regime shift. We compared the performance of the DL algorithm against lag-1 AC and variance for all six study systems.

## Results

The early warning signals provided by lag-1 AC, variance, and the DL algorithm can be compared as progressively more of the time series leading up to the bifurcation is made available for their computation, as might occur in real-world settings where a variable is monitored over time. For the two ecological models exhibiting the fold, Hopf and transcritical bifurcations (Figure 1a-c), the lag-1 AC and variance increase progressively before all three transition types except for the Hopf bifurcation, where lag-1 AC decreases due to the presence of an oscillatory component to the motion^18^) (Figure 1d-i). Hence the trends in these two indicators suggest that a transition will occur.

**Figure 1.**
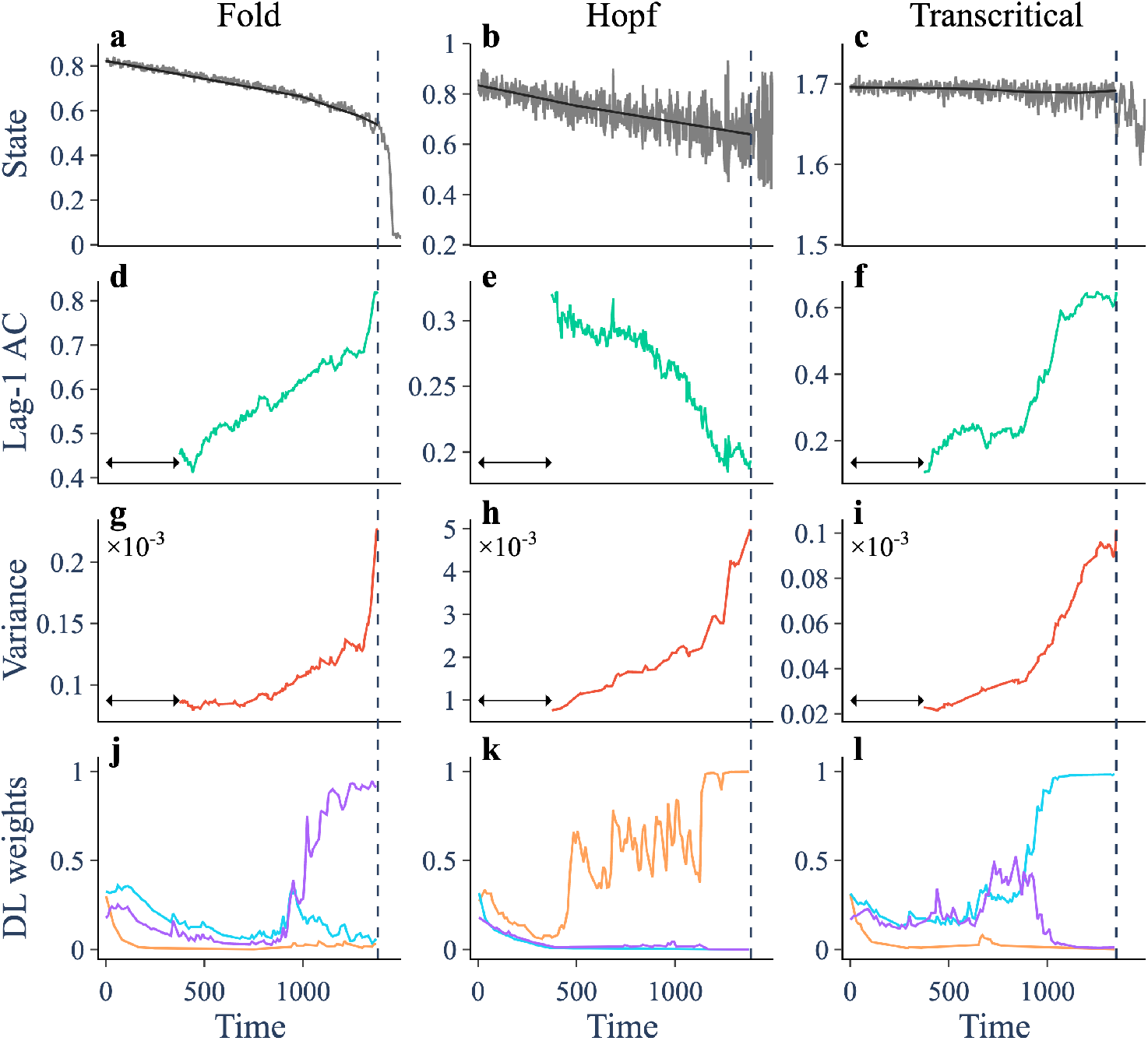
Trends in indicators prior to three different bifurcations in ecological models. (**a-c**) Trajectory (grey) and smoothing (black) of a simulation of an ecological model going through a fold, Hopf and transcritical bifurcation, respectively. (**d-f**) Lag-1 autocorrelation computed over a rolling window (arrow) of width 0.25. (**g-i**) Variance. (**j-l**) Weights assigned to the fold (purple), Hopf (orange) and transcritical (cyan) bifurcation by the DL algorithm.

The machine learning algorithm also indicates a transition, and correctly predicts the type of bifurcation in each of the three cases (Figure 1j-l). Inspection of the time series provides supporting evidence for our first two hypotheses. Firstly, these model equations were not used to develop our training library (although we note that our hypothesis relies on the training library including a representative type of dynamics from the models, such as Fold bifurcations). Secondly, the algorithm initially assigns similar probabilities to all three transition types in the earlier part of the time series, but after a specific time point, the algorithm becomes highly confident in picking one of the three bifurcation types as the most probable outcome. This is consistent with the algorithm being able to distinguish features based on higher-order terms that are held common between dynamical systems exhibiting each bifurcation type, but that distinguish the bifurcation types from one another. Examples of these time series for the other five study systems appear in the Appendix (Supplementary Figures S1-S10).

These time series figures, however, do not address how the approaches might perform when faced with a neutral time series where no transition occurs, and whether they might mistakenly generate a false positive prediction of an oncoming regime shift^29^. Hence, we compared the performance of these approaches with respect to both true and false positive through a receiver operator characteristics (ROC) curve. The ROC curve shows the ratio of true positives to false positives as a discriminant value that determines whether the classifier predicts a given outcome (such as transition versus no transition) is varied. The area under the ROC curve determines how well the classifier does with respect to both sensitivity (whether true positives are detected) and specificity (whether false positive are kept to a minimum). The AUC is 1 for a perfect classifier, and 0.5 for a classifier that is no better than random.

We compared ROC curves for the criterion of predicting any transition for lag-1 AC, variance and the DL algorithm, for eight comparisons across all six study systems (Figure 2). In support of our second hypothesis, these show that the DL algorithm strongly outperforms lag-1 AC and variance in six of the comparisons, although there are two interesting comparisons where the performance of the DL algorithm is similar to that of lag-1 AC or variance. For the SEIRx coupled behaviour-disease model, all of the classifiers are little better than random in the model variable for the number of infectious persons (*I*, Figure 2e). This occurs due to non-normality of the system associated with differing timescales for demographic and epidemiological processes^30^. However, the early warning signal is apparent in the variable *x* for the prevalence of pro-vaccine opinion, and the DL algorithm outperforms both lag-1 AC and variance in this respect. Also, for the paleo-climate data, the DL algorithm performs about as well as lag-1 AC, and both perform better than variance (2h). This may occur because the variance actually decreases before the transition in several of the empirical time series; the sampling data does not have high enough resolution; or because the system was forced too quickly.

**Figure 2.**
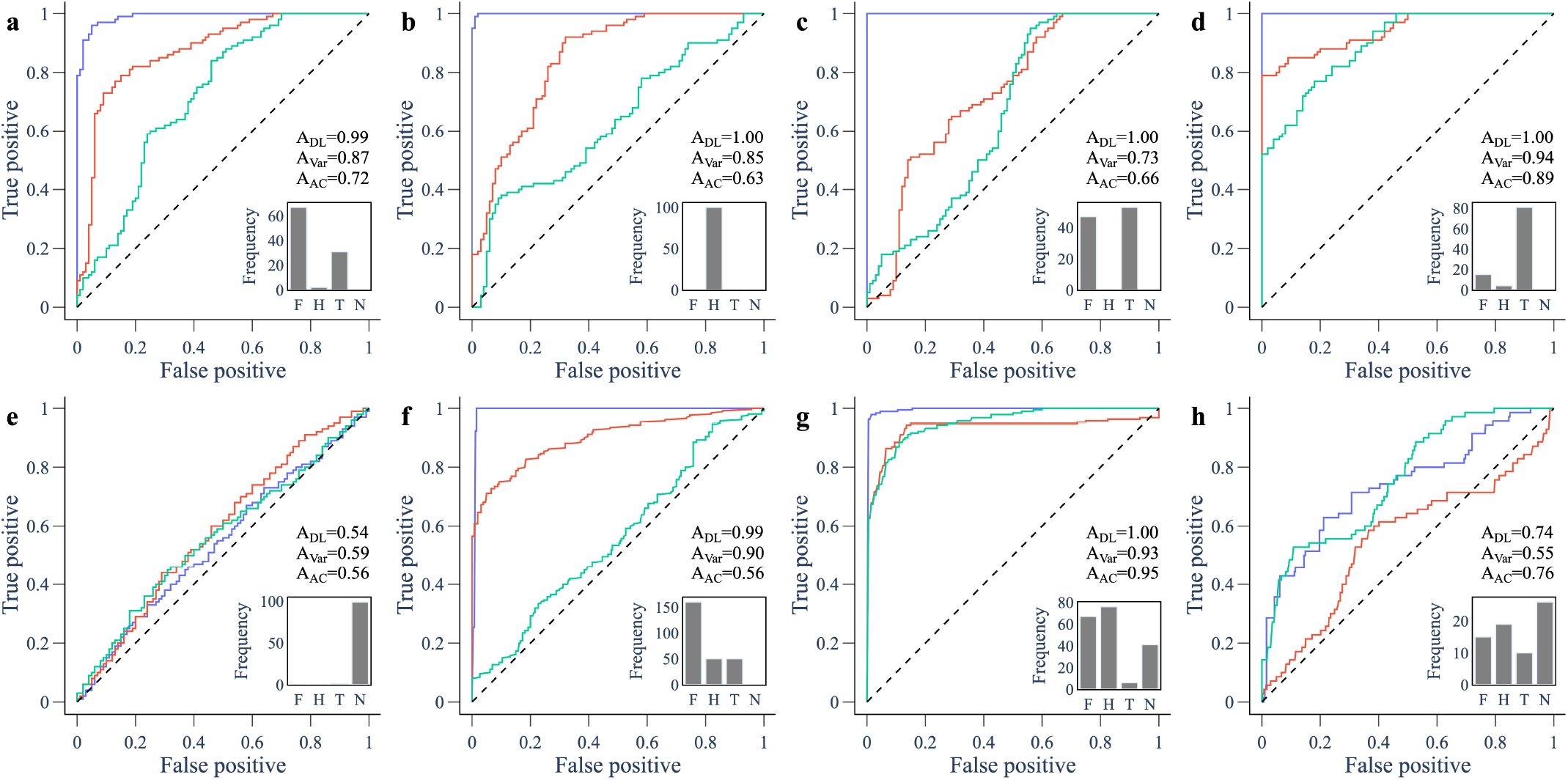
ROC curves for predictions using 80% – 100% of the pre-transition time series for model and empirical data. ROC curves compare the performance of the ML algorithm (blue), variance (red) and lag-1 autocorrelation (green) in predicting an upcoming transition. The area under the curve (AUC), abbreviated to A, is a measure of performance. The inset shows the frequency of the favoured ML weight among the forced trajectories: (F)old, (T)ranscritical, (H)opf, or (N)eutral. Panels show (**a**) May’s harvesting model going through a fold bifurcation; consumer-resource model going through a (**b**) Hopf and (**c**) transcritical bifurcation; behaviour-disease model going through a transcritical bifurcation using data from (**d**) pro-vaccine opinion (*x*) and (**e**) total infectious (*I*); (**f**) sediment data showing rapid transitions to an anoxic states in the Mediterranean sea; (**g**) data of a thermoacoustic system undergoing a Hopf bifurcation; (**h**) ice core records showing rapid transitions in paleoclimate data.

In support of our third hypothesis, we note that the DL classifier usually predicts the correct type of transition in all eight comparisons, but not always 2). For instance, the frequency of the favoured ML weight for Hopf bifurcation in the thermoacoustic system (Figure 2g) is only slightly higher than for the fold bifurcation. This could be due to our down sampling of the data to enable the time series to be accommodated by our classifier.

These ROC curves came from classifiers with access to 80% - 100% % of the time series (in other words, using the last 20% of the time series). We also computed the ROC curves when the classifiers have access to 60% - 80% (the fourth quintile) of the time series (Supplementary Appendix, Figure S11). This allows us to assess the reliability of the approaches when they are required to provide early warning when the system is still far from the regime shift. We observe that the DL algorithm provides early warning with greater sensitivity and specificity than either lag-1 AC or variance, in all comparisons except the *I* variable of the SEIRx model, where they perform equally poorly. This result suggests that the DL algorithm can provide greater forewarning of coming regime shifts, although additional statistical tests would be required to show this conclusively.

## Discussion

We tested our DL algorithm on data from systems that exhibited critical slowing down. However, other types of transitions are possible, such as global bifurcations that do not depend on changes to the local stability of fixed points^31^, or codimension two bifurcations that occur as two forcing parameters are varied simultaneously^14,32,33^. These regime shifts are not preceded by easily identified early warning signals^34^. In general, deep learning algorithms only work effectively for the specific problems they are trained to do. Therefore, in order for our algorithm to predict regime shifts for such bifurcations, we speculate that the training set would need to include those other bifurcation types. Similarly, we generated a training set based on models with two state variables and second-order polynomial model equations. This limits its ability to detect features such as deterministic chaos^20^, which require at least three state variables^35^.

Early warning indicators generally require high-resolution time series from a sufficiently long time series leading up to the tipping point^36^. This applies both to DL algorithms as well as lag-1 AC and variance. We did not analyze how the performance of the DL algorithms, lag-1 AC and variance compare as the time series becomes shorter. Similarly, none of these approaches can predict exactly when a transition will occur. This task lies in the domain of time series forecasting instead of classification and is a difficult undertaking for many systems, given that stochasticity could cause a system to jump prematurely to a new basin of attraction, even before a system has reached the tipping point^37^.

Other early warning approaches have been developed to predict the type of regime shift^18,20,30^. However, these approaches tend to be system-specific. Our results show that deep learning algorithms can improve not only the specificity and sensitivity of early warning signals for regime shifts, but also apply with a great degree of generality across different systems. Moreover, as long as the generic dynamical features (the normal forms) of the system near a tipping point are represented in the training set (Boxes 1,2), data from the study system are not required to train the algorithm. By combining dynamical systems insights with deep learning approaches, our results show how to predict and classify impending regime shifts with much greater sensitivity, specificity, and generalisability across systems than is currently possible. The type of regime shift is important to know theoretically, but also for practical purposes, since tipping points in many systems can lead to undesirable collapse^12^. Improved early warning signals will help us avoid and manage these regime shifts.

## Methods

### Generation of training data for the DL classifier

Training data consists of simulations of randomly generated, two-dimensional dynamical systems of the form

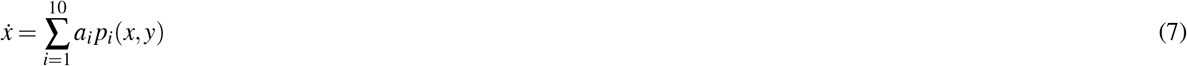

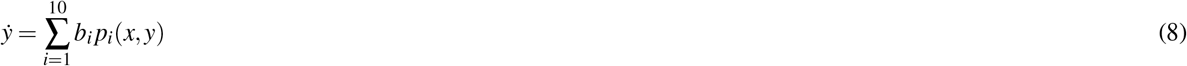

where *x* and *y* are state variables, *a*_*i*_ and *b*_*i*_ are parameters, and 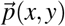 is a vector containing all polynomials in *x* and *y* up to third order

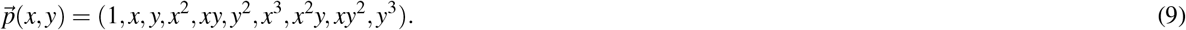

An individual model is generated by drawing each *a*_*i*_ and *b*_*i*_ from a normal distribution with zero mean and unit variance. Then, half of these parameters are selected at random and set to zero. The parameters for the cubic terms are set to the negative of their absolute value to encourage models with bounded solutions.

Upon generation of a model, we simulate it for 10,000 time steps from a randomly drawn initial condition and test for convergence to an equilibrium point. The simulation uses the odeint function from the Python package Scipy^38^ with a step size of 0.01. We say the model has converged if the maximum difference between the final 10 points of the simulation is less than 10^−8^. Models that do not converge are discarded. For models that converge, we use AUTO-07P^39^ to identify bifurcations along the equilibrium branch as each non-zero parameter is varied within the interval [-5, 5]. For each bifurcation identified, we run a corresponding ‘null’ and a ‘forced’ stochastic simulation of the model with additive white noise. Null simulations keep all parameters fixed. Forced simulations increase the bifurcation parameter linearly in time from its original value up to the bifurcation point. Stochastic simulations are run using the Euler Maruyama method with a step size of 0.01, an initial condition given by the models equilibrium value and a burn-in period of 100 units of time. The noise amplitude is drawn from a triangular distribution centred at 0.01 with upper and lower bounds 0.0125 and 0.0075 respectively, and weighted by an approximation of the dominant eigenvalue of the model (Supplementary Note).

For each simulation, we set a sampling rate *f*_*s*_, the number of data points collected per unit of time, which is drawn randomly from {1, 2, …, 10}. Using a varied sampling rate provided a wider distribution of lag-1 autocorrelation among the training data entries, which is important for representing a wide range of systems and timescales. The simulation is then run for 700*/ f*_*s*_ time units for the 500-classifier and 1700*/ f*_*s*_ time units for the 1500-classifier, providing 700 and 1700 points respectively when sampled at a frequency *f*_*s*_.

Due to noise, the simulations often transition to a new regime before bifurcation point is reached. We only want the DL classifier to see data prior to the transition. Therefore we use a change-point detection algorithm contained in the Python package *ruptures*^40^ to locate a transition point if one exists. If a transition point is detected, the preceding 500 (1500) points are taken as training data. If the transition occurs earlier than 500 (1500) data points into the simulation, the model is discarded. If no transition point is detected, the final 500 (1500) points are taken as training data.

We generated two different training sets: one consisting of 500, 000 time series of length 500 data points, and one consisting of 200, 000 time series of length 1500 data points. This was done because the time series lengths in the three empirical and three model systems are highly variable. The algorithm was trained separately on these two training sets (see next subsection), resulting in a “500-classifier” and a “1500-classifier”. The 500-classifier was used on shorter time series while the 1500-classifier was used on the longer time series. For the model time series in Figure 1, we use the 1500-classifier. For the ROC curves in Figure 2, we used the 500-classifier for the paleo-climate data and the ecological models; and the 1500-classifier for the thermoacoustic data, anoxia data, and disease model.

### Deep learning algorithm architecture and training

We used a convolutional neural network-long short-term memory (CNN-LSTM) deep learning algorithm^23,24^. We also experimented with a residual network, functional convolutional network, and recurrent neural network but found that the CNN-LSTM architecture yielded highest precision and recall on our training set. The code was written using TensorFlow 2.0 in Anaconda 2020.02. The CNN-LSTM architecture appears in Figure 3. The algorithm was trained for 1500 epochs with a learning rate of 0.0005, and the hyper-parameters were tuned through a series of grid sweeps. The same hyper-parameter values were used for training on both 500-classifier and 1500-classifier (see previous subsection).

**Figure 3.**
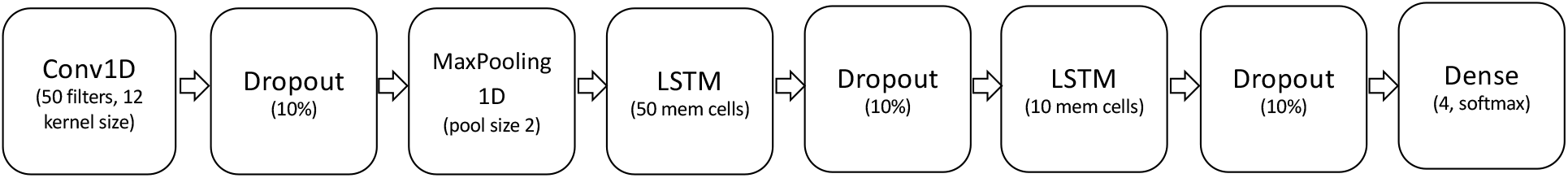
CNN-LSTM architecture.

The simulation output time series from the random dynamical systems were detrended using Lowess smoothing with a span of 0.2 to obtain the residual time series that formed the training set. Each residual time series was normalized by dividing each time series datapoint by the average absolute value of the residuals across the entire time series. We used a train/validation/test split of 0.95*/*0.04*/*0.01 for both the 500- and 1500-classifiers. The test set was chosen as a small percentage because a test set of a few thousand time series is adequate to provide a representative estimate of the precision and recall. The f1 score, precision and recall for an ensemble of ten 500-classifier models were 84.2%, 84.4%, and 84.2%, respectively. The f1 score, precision and recall for an ensemble of ten 1500-classifier models were 88.2%, 88.3%, and 88.3%, respectively.

For testing the ability of the DL algorithm to provide early warning signals of bifurcations, we developed variants where the algorithm was trained on censored versions of the training time series. For the 500 (1500) length classifier, one variant was trained on a version of the training set where the residuals of the simulation time series were padded on both the left and right by between 0 and 225 (725) zeroes, with the padding length chosen randomly from a uniform distribution. This allowed the algorithm to train on time series as short as 50 (50) not necessarily representing the time phase just before the transition. The intention was to boost the performance of the DL algorithm for detecting EWS features from shorter time series and from the middle sections of time series. The second variant was trained on a version of the training set where the residuals of the simulation time series were padded only on the left, by between 0 and 450 (1450) zeroes where the padding length was chosen randomly from a uniform distribution. This allowed the algorithm to train on time series as short as 50 (50) representing time series of various lengths that lead up to the bifurcation (except for the neutral class). As a result, the classifier could better detect features that emerge most strongly right before the bifurcation. Ten trained models of each variant were ensembled by taking their average prediction at each point to generate all of our reported results.

### Theoretical models used for testing

We use models of low and intermediate complexity to test the DL classifier. Models are simulated using the Euler Maruyama method with a step size of 0.01 unless otherwise stated. To test detection of a Fold bifurcation, we use May’s harvesting model^25^ with additive white noise. This is given by

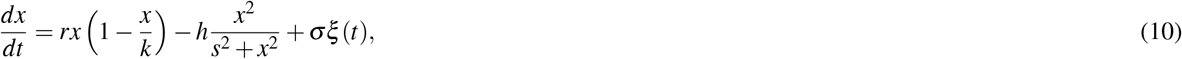

where *x* is biomass of some population, *k* is its carrying capacity, *h* is the harvesting rate, *s* characterises the nonlinear dependence of harvesting output on current biomass, *σ* is the noise amplitude, and *ξ* (*t*) is a Gaussian white noise process. We use parameter values *r* = 1, *k* = 1, *s* = 0.1, *h* ∈ [0.15, 0.27] and *σ* = 0.01. In this configuration, a Fold bifurcation occurs at *h* = 0.26. The parameter *h* is kept fixed at its lower bound for null simulations and is increased linearly to its upper bound in forced simulations.

To test the Hopf and Transcritical bifurcations, we use the Rozenzweig-MacArthur consumer-resource model^26^ with additive white noise. This is given by

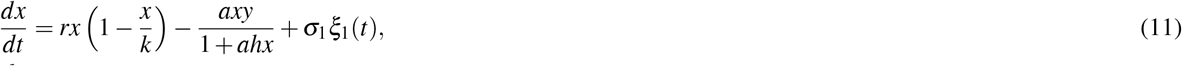

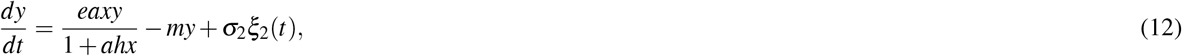

where *r* is the intrinsic growth rate of the resource (*x*), *k* is its carrying capacity, *a* is the attack rate of the consumer (*y*), *e* is the conversion factor, *h* is the handling time, *m* is the consumer mortality rate, *σ*_1_ and *σ*_2_ are noise amplitudes, and *ξ*_1_(*t*) and *ξ*_2_(*t*) are independent Gaussian white noise processes. We fix the parameter values *r* = 4, *k* = 1.7, *e* = 0.5, *h* = 0.15, *m* = 2, *σ*_1_ = 0.01, and *σ*_2_ = 0.01. In this configuration, the deterministic system has a Transcritical bifurcation at *a* = 5.60 and a Hopf bifurcation at *a* = 15.69. For the Transcritical bifurcation, we simulate null trajectories with *a* = 2 and forced trajectories with *a* ∈ [2, 6]. For the Hopf bifurcation, we use *a* = 12 and *a* ∈ [12, 16] respectively.

To test the DL algorithm on a model of higher dimensionality than the ecological models, we used a stochastic version of the SEIRx model that captures interactions between disease dynamics and population vaccinating behaviour^11,41^ given by

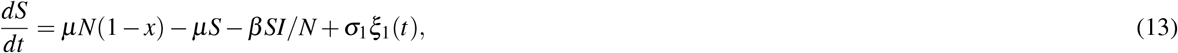

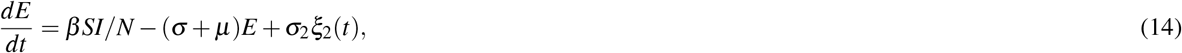

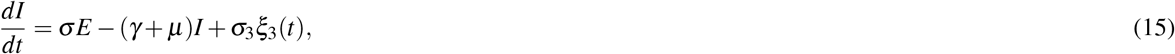

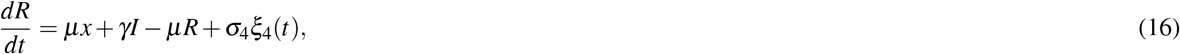

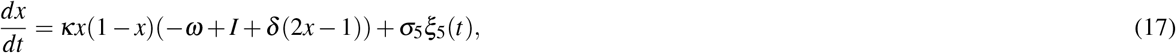

where *S* is the number of susceptible individuals, *E* is the number of exposed (infected but not yet infectious) individuals, *I* is the number of infectious individuals, *R* is the number of recovered/immune individuals, *x* is the number of individuals with pro-vaccine sentiment, *µ* is the per capita birth and death rate, *β* is the transmission rate, *σ* is the rate at which exposed individuals become infectious, *γ* is the rate of recovery from infection, *κ* is the social learning rate, *δ* is the strength of injunctive social norms, and *ω* is the perceived relative risk of vaccination versus infection. For our simulations we used *µ* = 0.02*/yr, β* = 1.5*/day* (based on *R*_0_ ≈ *β/γ* = 15), *σ* = 0.1*/day, γ* = 0.1*/day, κ* = 0.001*/day, δ* = 50, *N* = 100, 000 representing a typical paediatric infectious disease^11^. Simulations were perturbed weekly with *σ*_*i*_ = 5 for *i* = 1 … 4, and *σ*_5_ = 5 × 10^−4^. On account of the large timescale difference in vital dynamics and infection processes, the systems is non-normal^30^. We note that *S* + *E* + *I* + *R* = 1 and therefore since *R* can be obtained as *R* = 1 − *S* − *E* − *I*, the model is four dimensional. The forcing parameter *ω* was gradually forced from 0 to 100. As perceived vaccine risk increases along the (1, 0, 0, 0, 1) branch corresponding to full vaccine coverage, the model has a transcritical bifurcation at *ω* = *δ* ^11^, which leads to a critical transition corresponding to a drop in the proportion of individuals with pro-vaccine sentiment and a return of endemic infection.

### Empirical systems used for testing

We use three different sources of empirical data to test the DL classifier.

#### 1. Sedimentary archives from the Mediterranean sea^28^

These provide high-resolution reconstructions of oxygen dynamics in the eastern Mediterranean sea. Rapid transitions between oxic and anoxic states occurred regularly in this region in the geological past. A recent study has shown that early warning signals exist prior to the transitions^28^. The data consists of output from three cores that together span 8 anoxic events. Variables include Molybdenum (Mo) and Uranium (U), proxies for anoxic and suboxic conditions, respectively, giving us a total of 26 time series for anoxic events (some are captured by multiple cores). The sampling rate provides *∼*10-50 year resolution depending on the core with an almost regular spacing between data points. We perform the same data preprocessing as Hennekam et al.^28^. Interpolation is not done, as most data points are equidistant and it can give rise to aliasing effects that strongly affect variance and autocorrelation. Data 10kyr prior to each transition are analysed for early warning signals. Null time series of the same length are generated from an AR(1) process fit to the initial 20% of the data. Residuals are obtained from smoothing the data with a Gaussian kernel with a bandwidth of 900yr and EWS are computed using a rolling window of 0.5. The geochemical data are available at the PANGEA repository https://doi.pangaea.de/10.1594/PANGAEA.923197.

#### 2. Thermoacoustic instability

Thermoacoustic systems often exhibit a critical transition to a state of self-sustained large amplitude oscillations in the system variables, known as thermoacoustic instability. The establishment of a positive feedback between the heat release rate fluctuations and the acoustic field in the system is often the cause for this transition. We perform experiments in a horizontal Rijke tube which consists of an electrically heated wire mesh in a rectangular duct^27^. We pass a constant mass flow rate of air through the duct and control the voltage applied across the wire mesh to attain the transition to thermoacoustic instability via subcritical Hopf bifurcation as the voltage is increased. We have data for 19 forced trajectories where the voltage is increased over time at different rates (2-24000 mV/s). We also have 10 steady state trajectories where the voltage is kept at a fixed value between 0 and 4V. Experimental runs at fixed higher voltages are not used as they exhibit limit cycle oscillations. We downsample the data from 4-10kHz in experiments to 2kHz. Transition times are picked by eye and provided in Supplementary Table. For each forced time series, we analyse data 1500 points prior to the transition. From the steady state time series, we extract two random sections of length 1500 to serve as null time series, giving a total of 20 null time series. Data is detrended using Lowess smoothing with a span of 0.2 and degree 1. EWS are computed from residuals using a rolling window of 0.5.

#### 3. Paleoclimate transitions

We use data for 7 out of the 8 climate transitions that were previously analysed for EWS by Dakos et al.^15^ Data are available from sources specified therein. Time series for the desertification of N. Africa was not included due to insufficient data. We use the same data preprocessing as Dakos et al.^15^, which involves using linear interpolation to make the data equidistant, and detrending with a Gaussian kernel smoothing function. Bandwidth of the kernel is specified for each time series^15^ to remove long-term trends whilst not overfitting. For each time series, we generate 10 null time series of the same length from an AR(1) process fit to the initial 20% of the residuals, yielding a total of 70 null time series and 7 forced time series.

### Computing early warning indicators and comparing predictions with the DL classifier

Generic early warning indicators are computed using the Python package *ewstools*^18^ which implements established methods^42^. This first involves detrending the time series to obtain residual dynamics. This is done using Lowess smoothing^43^ with span 0.2 and degree 1 unless stated otherwise. Variance and lag-1 autocorrelation are then computed over a rolling window of length To assess the presence of an early warning signal, we use the Kendall *τ* value, which serves as a measure of increasing or decreasing trend. The Kendall tau value at a given time is computed over the entire preceding data.

To compare predictions made between the conventional EWS and the DL classifier, we use receiver operator characteristics (ROC). The ROC curve shows the ratio of true positives to false positives as a discriminant value is varied. For the EWS, the discriminant value is the Kendall tau value. For the DL classifier, the discriminant value is the weight given to a bifurcation trajectory.

## Acknowledgements

We thank Ryan Kinnear (University of Waterloo) for helpful discussions on time series analysis, and Rick Hennekam and Gert-Jan Reichart for providing the anoxia dataset.

## Supplementary Information

## 1 Supplementary Figures

**Supplementary Figure 1.**
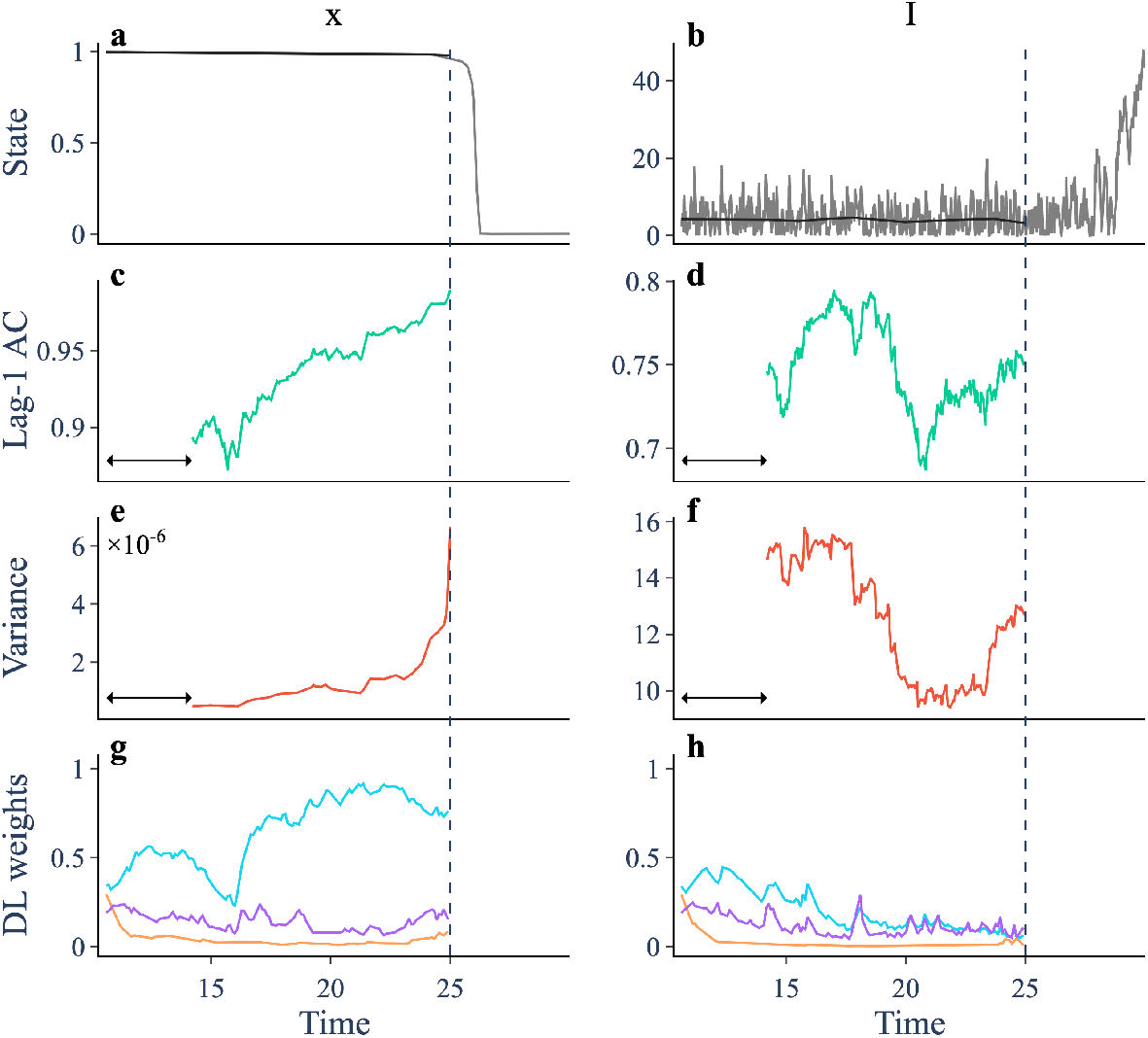
Early warning signals and DL predictions prior to a vaccine scare in the SEIRx model. (a-b) Trajectory (grey) and smoothing (black) of a model simulation showing the vaccine uptake (x) and the proportion infected (I). The dashed vertical line identifies the onset of the transition. (d-f) Lag-1 autocorrelation computed over a rolling window (arrow) of size 0.25. (g-i) Variance. (j-l) ML weights for a fold (purple), Hopf (orange) and transcritical (cyan) bifurcation.

**Supplementary Figure 2.**
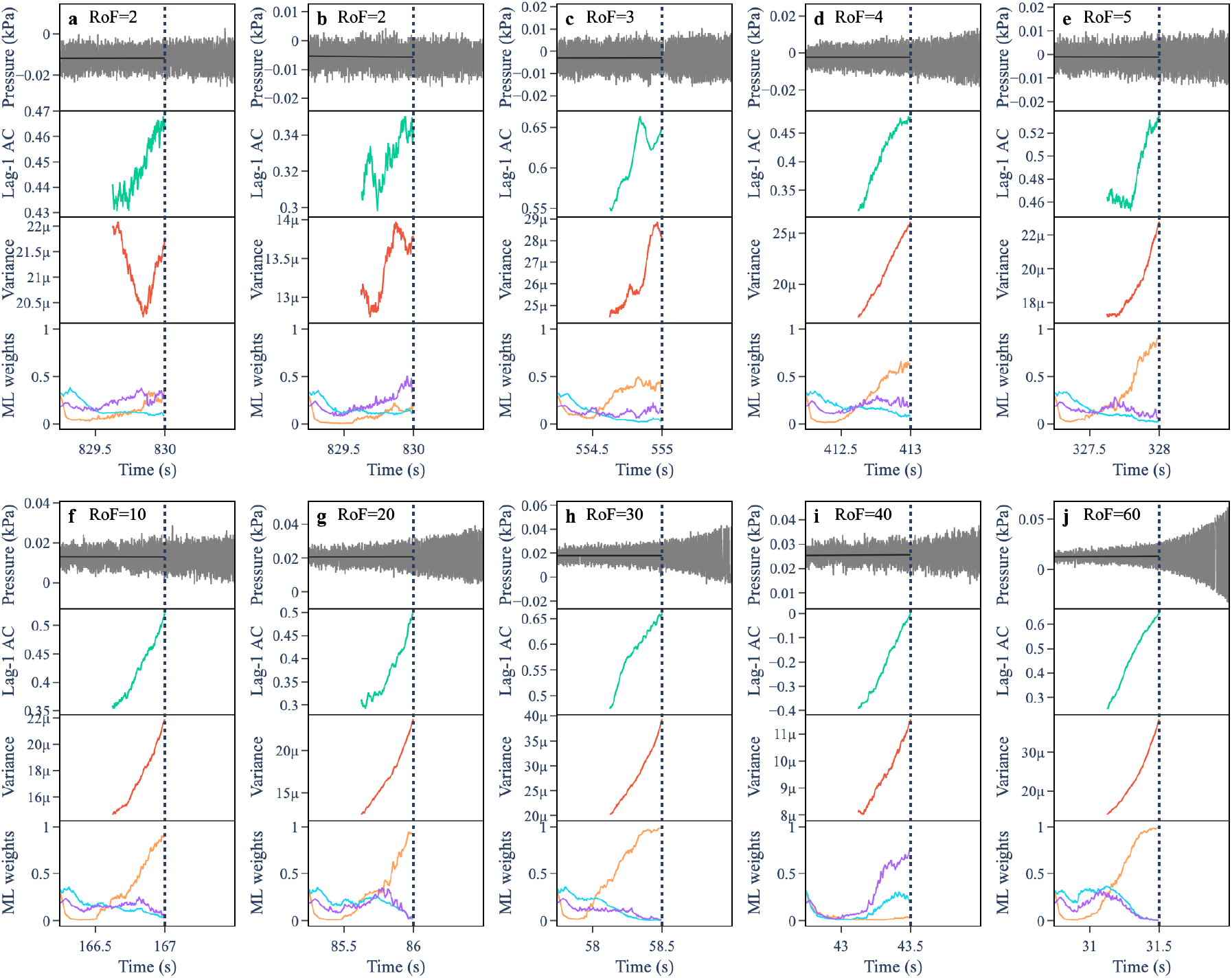
Early warning signals and DL predictions for thermoacoustic transitions. (a-j) Each panel shows a thermoacoustic transition and leading indicators for a given rate of forcing (RoF). (Top) Trajectory (grey) and smoothing (black). (2nd down) Lag-1 autocorrelation computed over a rolling window of length 0.5. (3rd down) Variance computed over a rolling window of length 0.5. (Bottom) ML weights for a fold (purple), Hopf (orange) and transcritical (cyan) bifurcation.

**Supplementary Figure 3.**
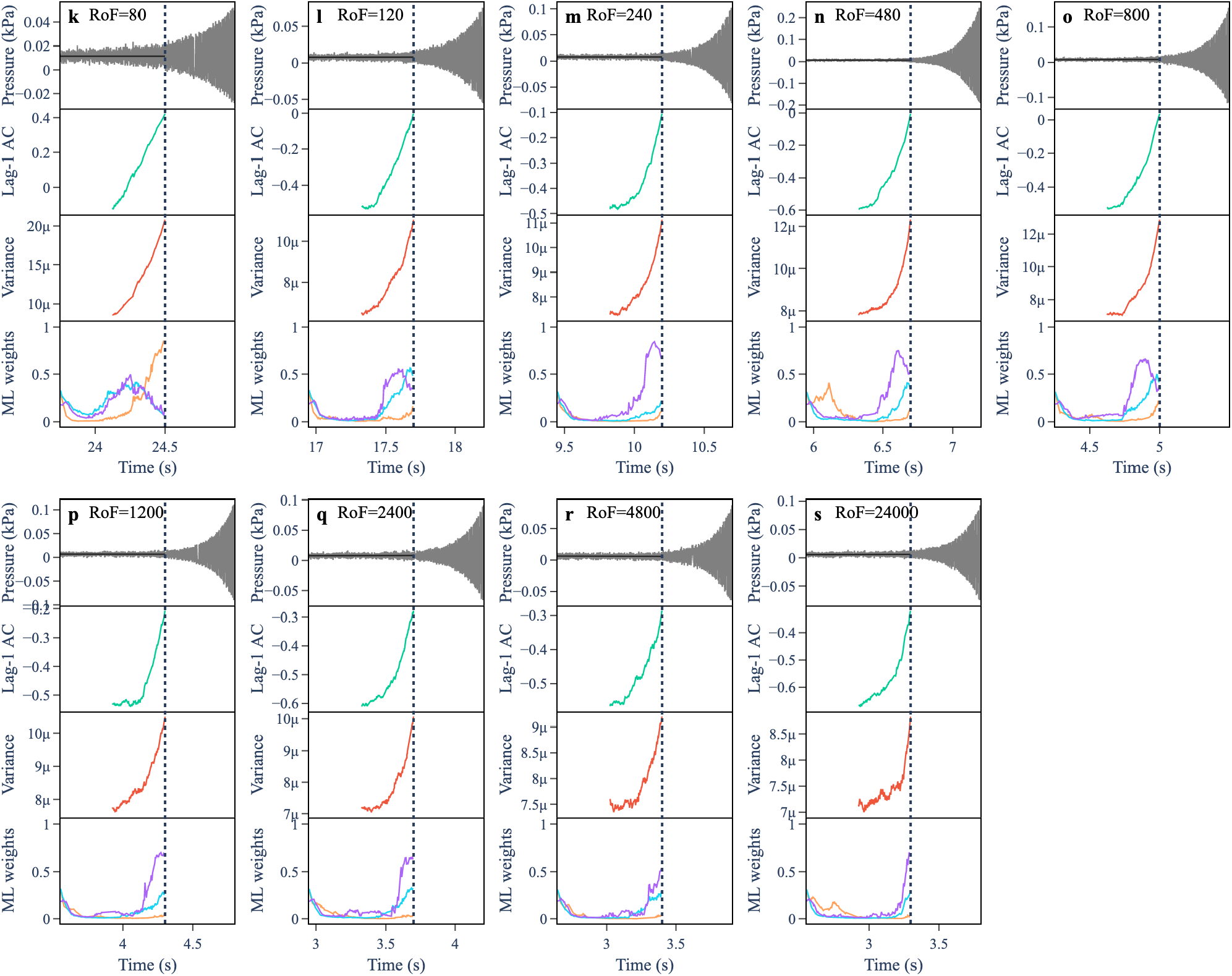
Early warning signals and DL predictions for thermoacoustic transitions (continued). (k-s) Each panel shows a thermoacoustic transition and leading indicators for a given rate of forcing (RoF). (Top) Trajectory (blue) and Lowess smoothing (grey). (2nd down) Lag-1 autocorrelation computed over a rolling window of length 0.5. (3rd down) Variance computed over a rolling window of length 0.5. (Bottom) ML weights for a fold (purple), Hopf (orange) and transcritical (cyan) bifurcation.

**Supplementary Figure 4.**
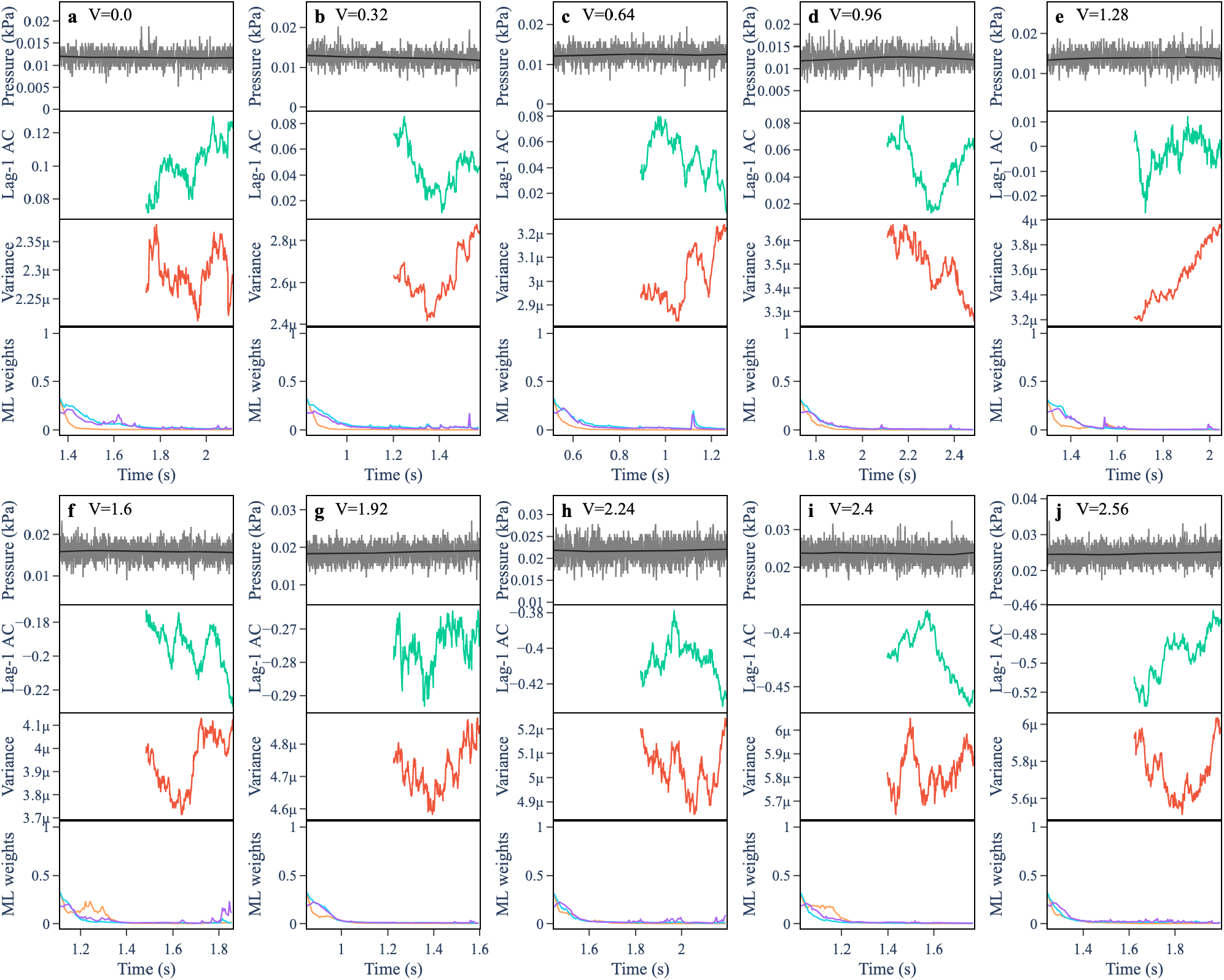
Early warning signals and DL predictions for null thermoacoustic trajectories. (a-j) Each panel shows a thermoacoustic trajectory at some fixed voltage (V). (Top) Trajectory (blue) and Lowess smoothing (grey). (2nd down) Lag-1 autocorrelation computed over a rolling window of length 0.5. (3rd down) Variance computed over a rolling window of length 0.5. (Bottom) ML weights for a fold (purple), Hopf (orange) and transcritical (cyan) bifurcation.

**Supplementary Figure 5.**
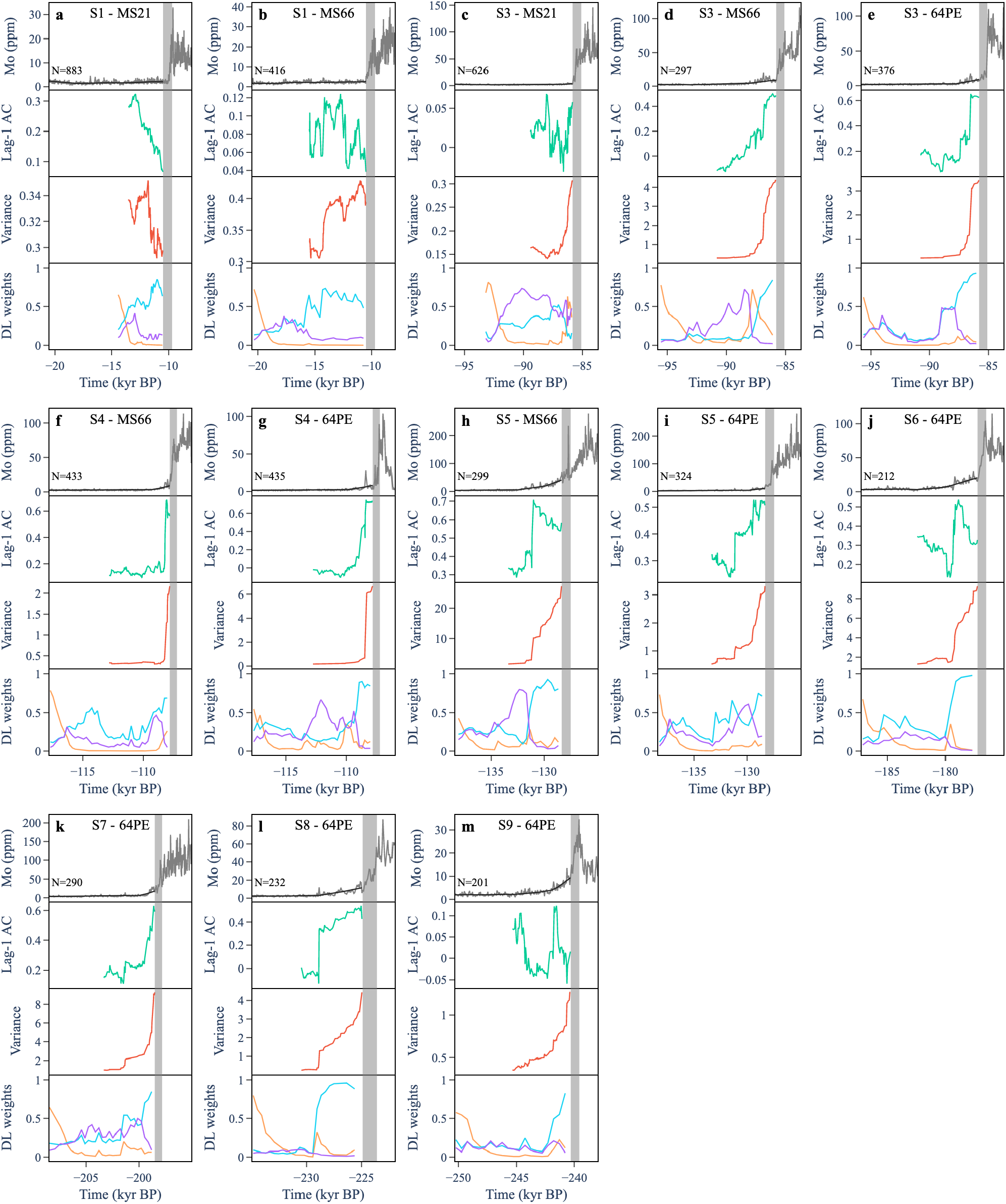
Early warning signals and DL predictions in Molybdenum (Mo) data for 8 anoxic transitions. (a-j) Each panel shows a past anoxic transition (S1-S9) from data in a given core (MS21, MS66, or 64PE406E1). (Top) Trajectory (blue) and Gaussian smoothing (grey). (2nd down) Lag-1 autocorrelation computed over a rolling window of length 0.5. (3rd down) Variance computed over a rolling window of length 0.5. (Bottom) ML weights for a fold (purple), Hopf (orange) and transcritical (cyan) bifurcation.

**Supplementary Figure 6.**
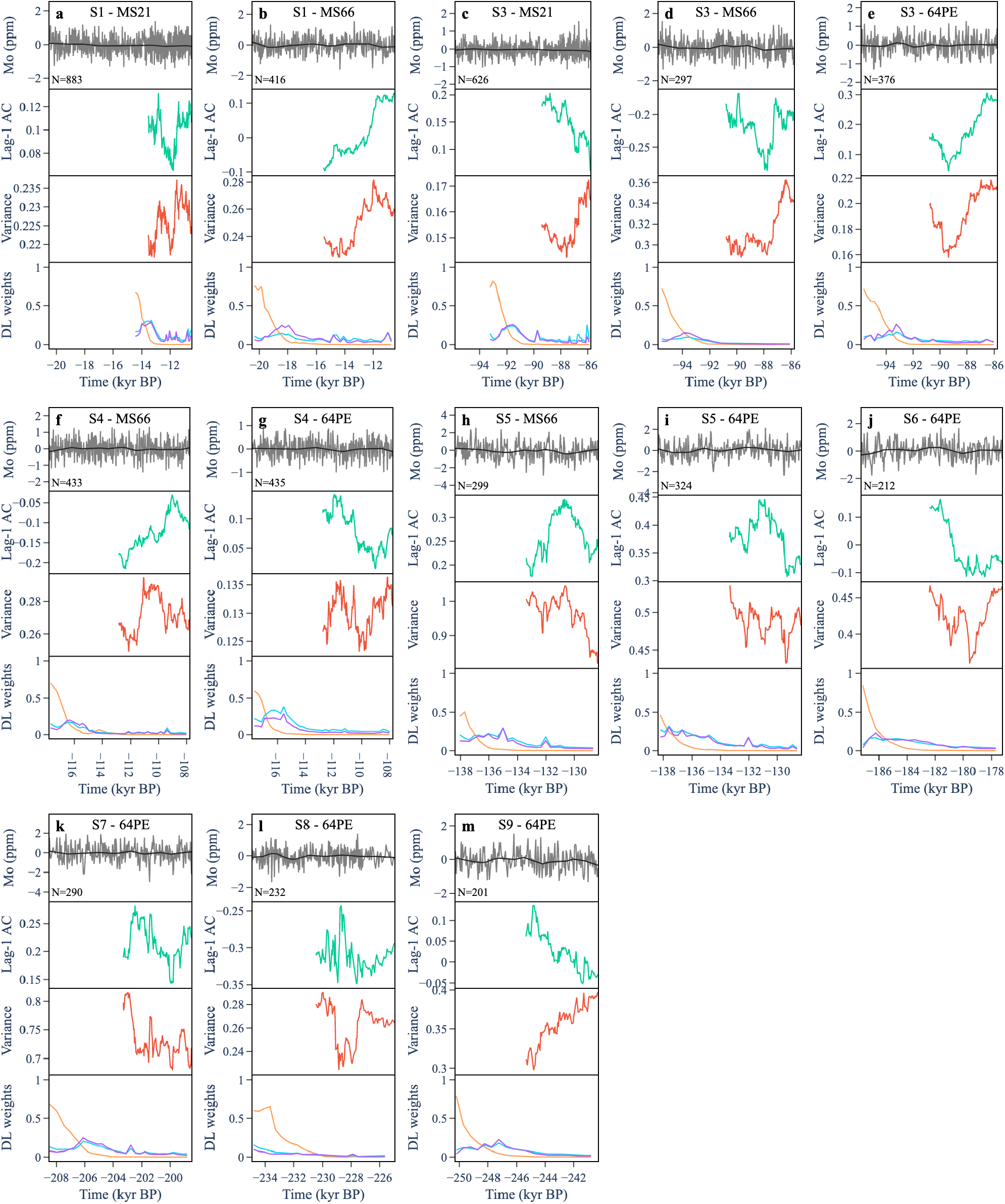
Early warning signals and DL predictions for simulated null time series of Molybdenum (Mo). (a-j) Each panel shows a simulated null trajectory from an AR1 process fit to the first 20% of anoxic transitions (S1-S9) from data in a given core (MS21, MS66, or 64PE406E1). (Top) Trajectory (blue) and Gaussian smoothing (grey). (2nd down) Lag-1 autocorrelation computed over a rolling window of length 0.5. (3rd down) Variance computed over a rolling window of length 0.5. (Bottom) ML weights for a fold (purple), Hopf (orange) and transcritical (cyan) bifurcation.

**Supplementary Figure 7.**
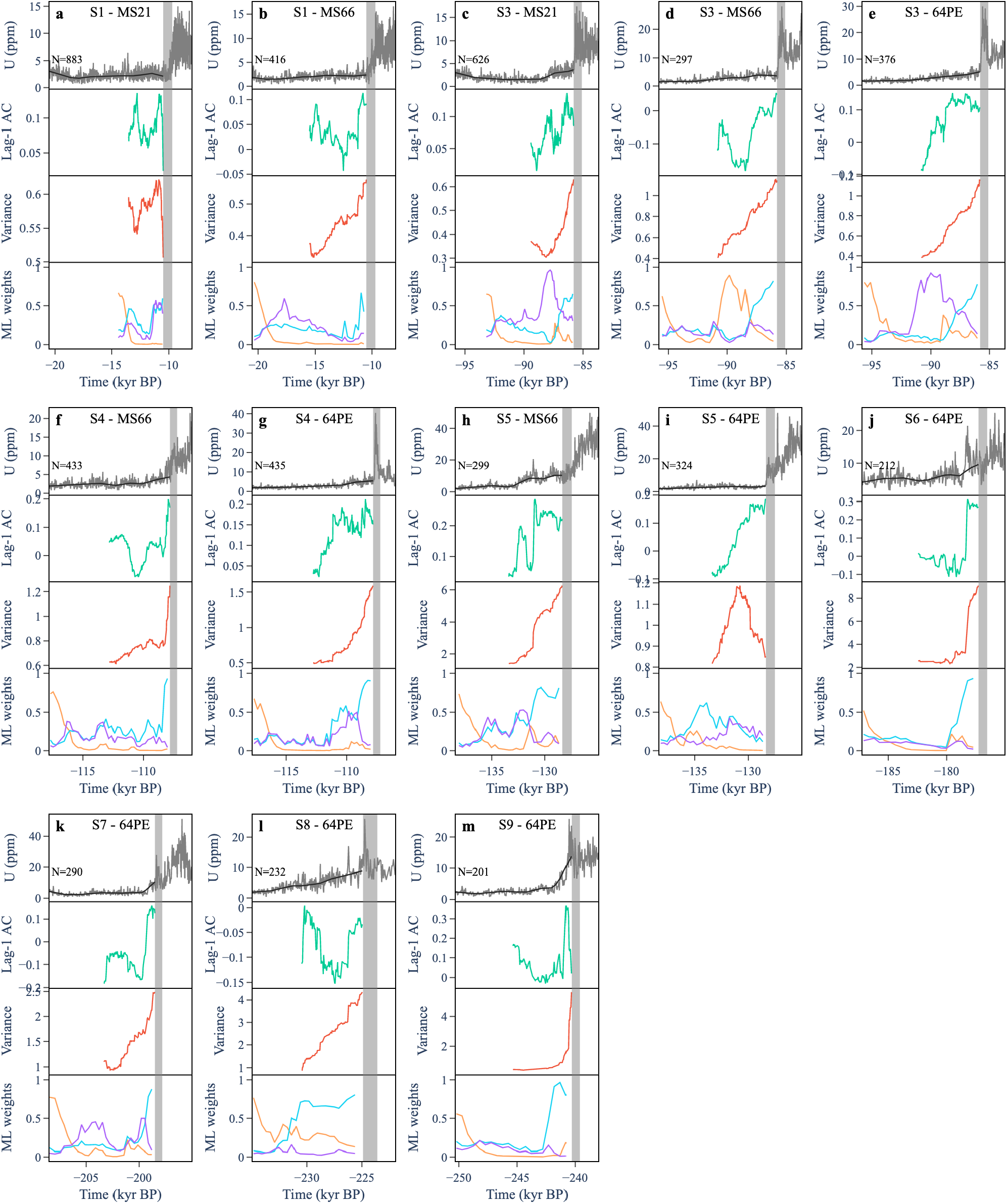
Early warning signals and DL predictions in Uranium (U) data for 8 anoxic transitions. (a-j) Each panel shows a past anoxic transition (S1-S9) from data in a given core (MS21, MS66, or 64PE406E1). (Top) Trajectory (blue) and Gaussian smoothing (grey). (2nd down) Lag-1 autocorrelation computed over a rolling window of length 0.5. (3rd down) Variance computed over a rolling window of length 0.5. (Bottom) ML weights for a fold (purple), Hopf (orange) and transcritical (cyan) bifurcation.

**Supplementary Figure 8.**
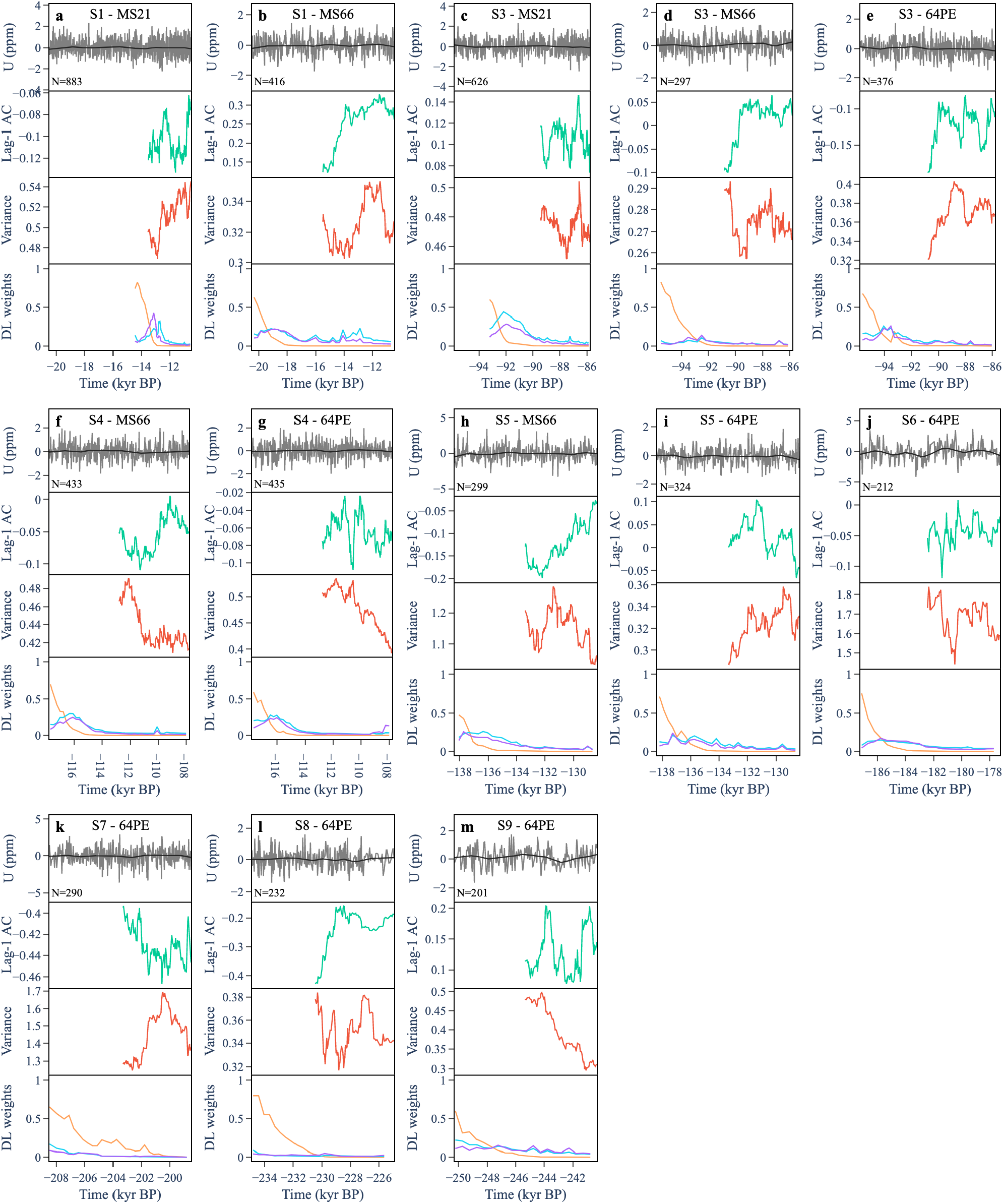
Early warning signals and DL predictions for simulated null time series of Uranium (U). (a-j) Each panel shows a simulated null trajectory from an AR1 process fit to the first 20% of anoxic transitions (S1-S9) from data in a given core (MS21, MS66, or 64PE406E1). (Top) Trajectory (blue) and Gaussian smoothing (grey). (2nd down) Lag-1 autocorrelation computed over a rolling window of length 0.5. (3rd down) Variance computed over a rolling window of length 0.5. (Bottom) ML weights for a fold (purple), Hopf (orange) and transcritical (cyan) bifurcation.

**Supplementary Figure 9.**
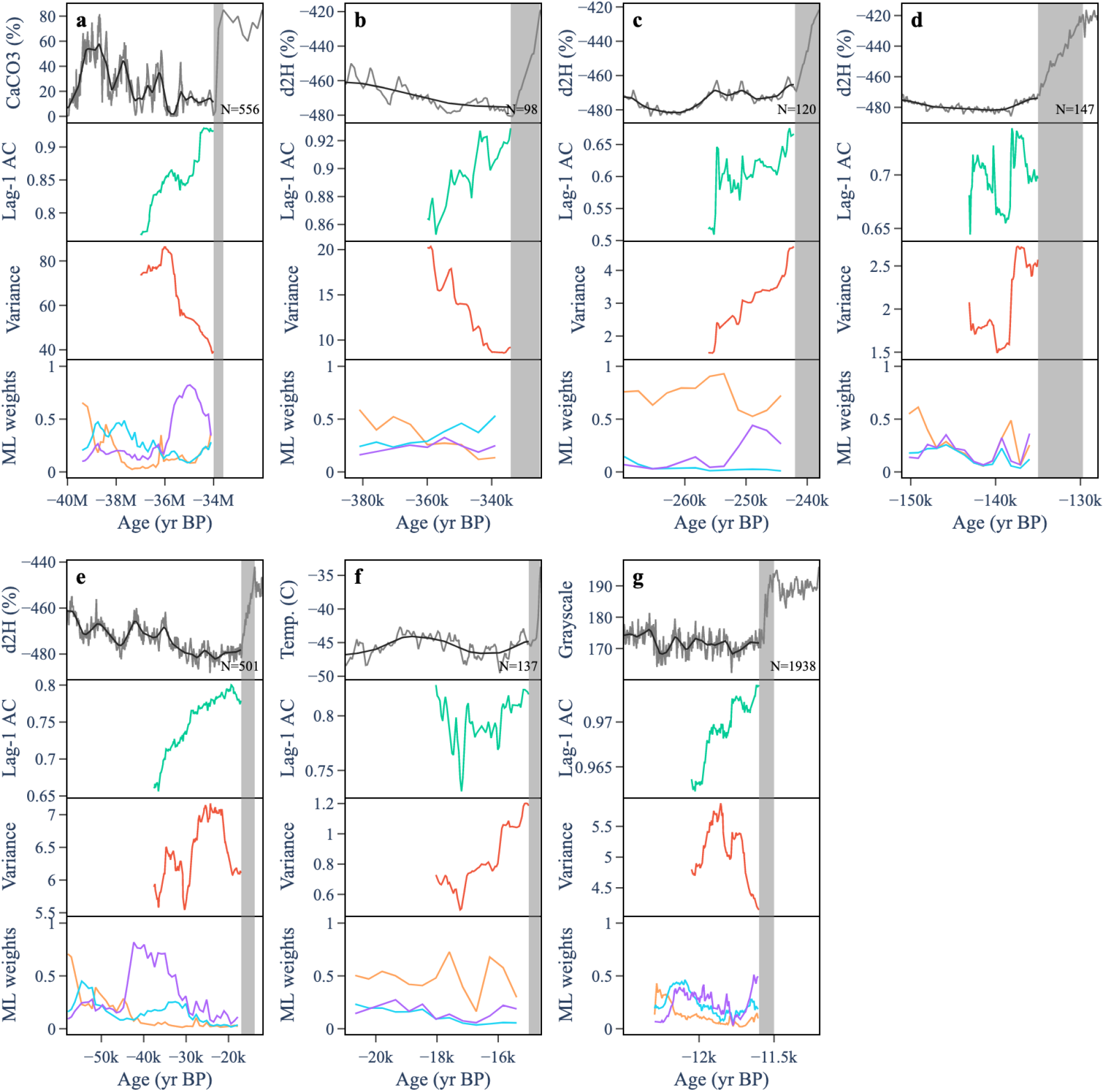
Early warning signals and DL predictions for paleoclimate transitions. a) End of greenhouse Earth, b) End of glaciation IV, c) End of glaciation III, d) End of glaciation II, e) End of glaciation I, f) Bolling/Allerod transition, g) End of Younger Dryas. Panels are organised chronologically from oldest to youngest. (Top) Trajectory (blue) and Gaussian smoothing (grey). (2nd down) Lag-1 autocorrelation computed over a rolling window of length 0.5. (3rd down) Variance computed over a rolling window of length 0.5. (Bottom) ML weights for a fold (purple), Hopf (orange) and transcritical (cyan) bifurcation.

**Supplementary Figure 10.**
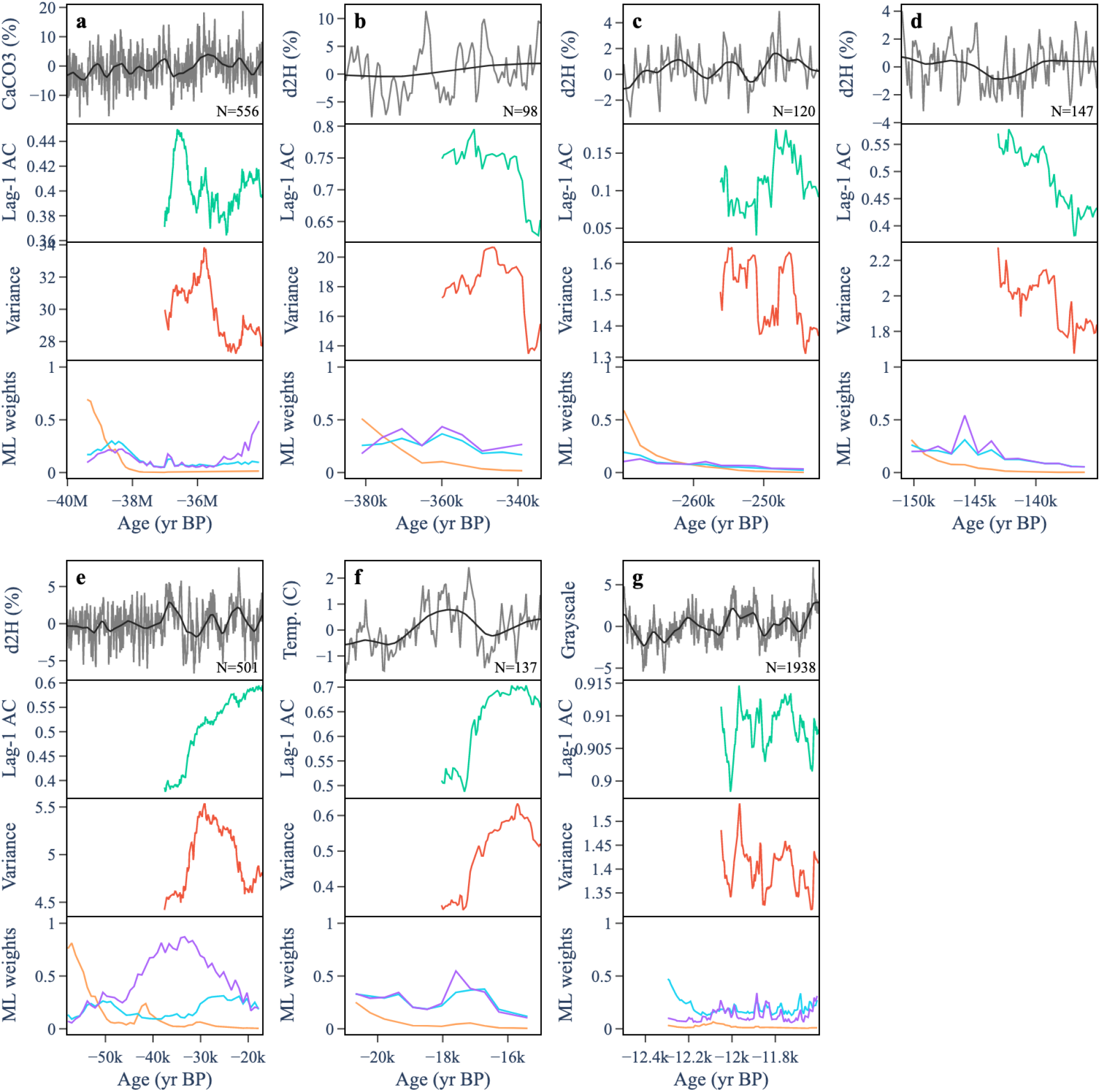
Early warning signals and DL predictions in simulated null trajectories for paleoclimate data. a) End of greenhouse Earth, b) End of glaciation IV, c) End of glaciation III, d) End of glaciation II, e) End of glaciation I, f) Bolling/Allerod transition, g) End of Younger Dryas. Panels are organised chronologically from oldest to youngest. (Top) Trajectory (blue) and Gaussian smoothing (grey). (2nd down) Lag-1 autocorrelation computed over a rolling window of length 0.5. (3rd down) Variance computed over a rolling window of length 0.5. (Bottom) ML weights for a fold (purple), Hopf (orange) and transcritical (cyan) bifurcation.

**Supplementary Figure 11.**
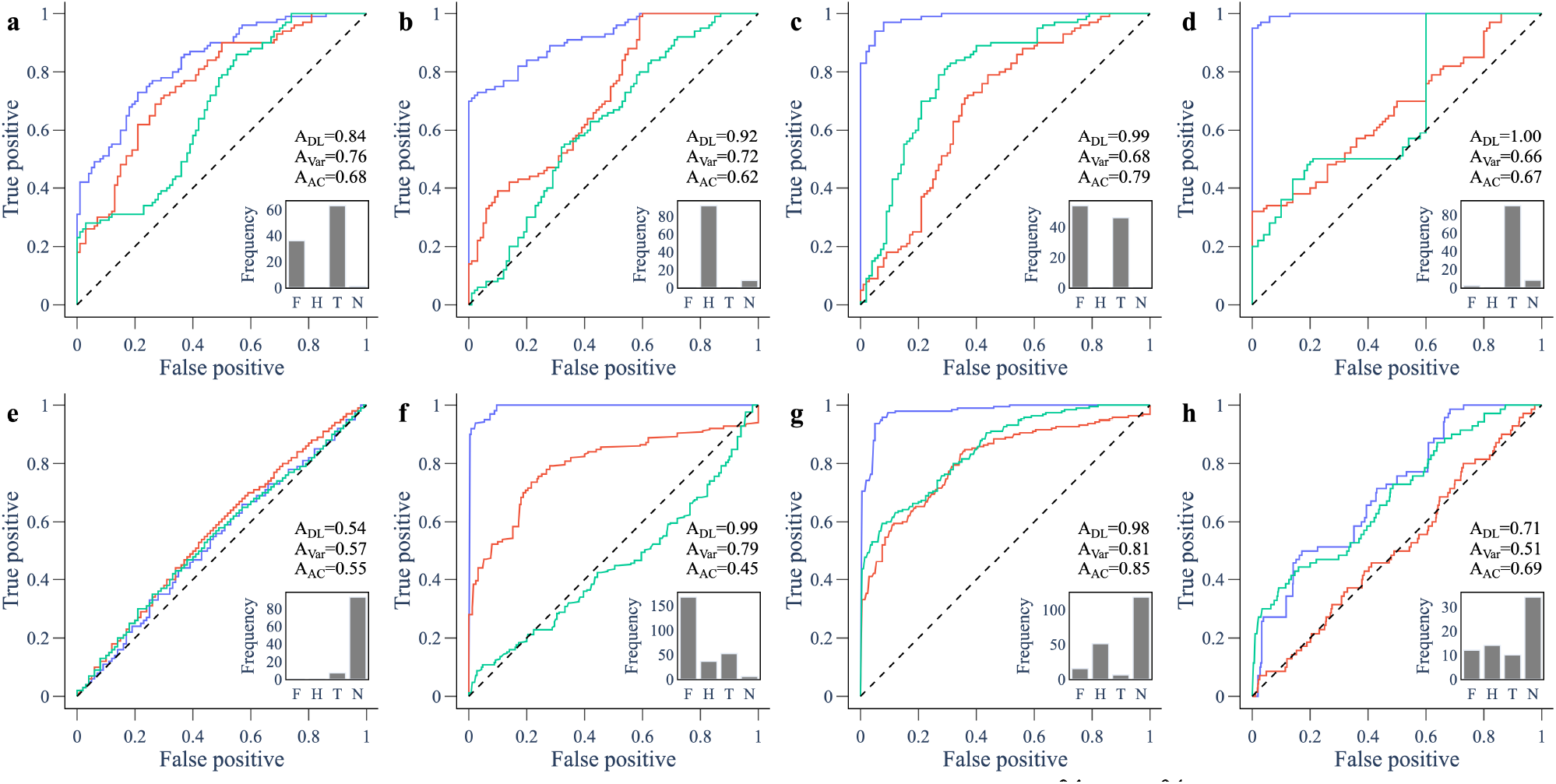
ROC curves for predictions using 60% – 80% of the pre-transition time series for model and empirical data. ROC curves compare the performance of the ML algorithm (blue), variance (red) and lag-1 autocorrelation (green) in predicting an upcoming transition. The area under the curve (AUC), abbreviated to A, is a measure of performance. The inset shows the frequency of the favoured ML weight among the forced trajectories: (F)old, (T)transcritical, (H)opf, or (N)eutral. Panels show (**a**) May’s harvesting model going through a fold bifurcation; consumer-resource model going through a (**b**) Hopf and (**c**) transcritical bifurcation; behaviour-disease model going through a transcritical bifurcation using data from (**d**) pro-vaccine opinion (*x*) and (**e**) total infectious (*I*); (**f**) sediment data showing rapid transitions to an anoxic states in the Mediterranean sea; (**g**) data of a thermoacoustic system undergoing a Hopf bifurcation; (**h**) ice core records showing rapid transitions in paleoclimate data.

## 2 Supplementary Note

### 2.1 Noise amplitude of simulations in training data

We would like to select a noise amplitude that lies in an appropriate range for each of the generated models. Ideally, simulations will have a noise amplitude that is large enough such that nonlinear terms are present in the signal, but small enough so that noise does not swamp the deterministic structure of the model. This amplitude is model-dependent, and can be approximated based on the size of the dominant eigenvalue of the generated model.

In a small neighbourhood of the equilibrium point, the dynamics of the training model are well described by its linearization

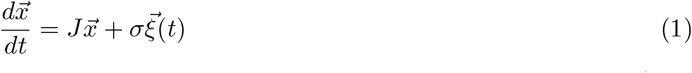

where 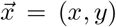 are the state variables, *J* is the Jacobian matrix, *σ* is the noise amplitude and 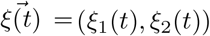 are the white noise processes. This equation describes an Ornstein-Uhlenbeck process, for which the variance can be computed analytically [1]. To first-approximation, this variance is given by

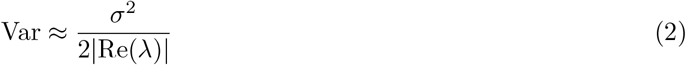

where *λ* is the dominant eigenvalue of *J*, that is, the eigenvalue whose real part has the smallest magnitude. The dominant eigenvalue can be computed for each dynamical system that we generate. This approximate relationship is shown by simulating an ensemble of models with fixed, small noise *σ* = 0.01, and plotting the variance against the size of the dominant eigenvalue (Figure 12).

**Supplementary Figure 12.**
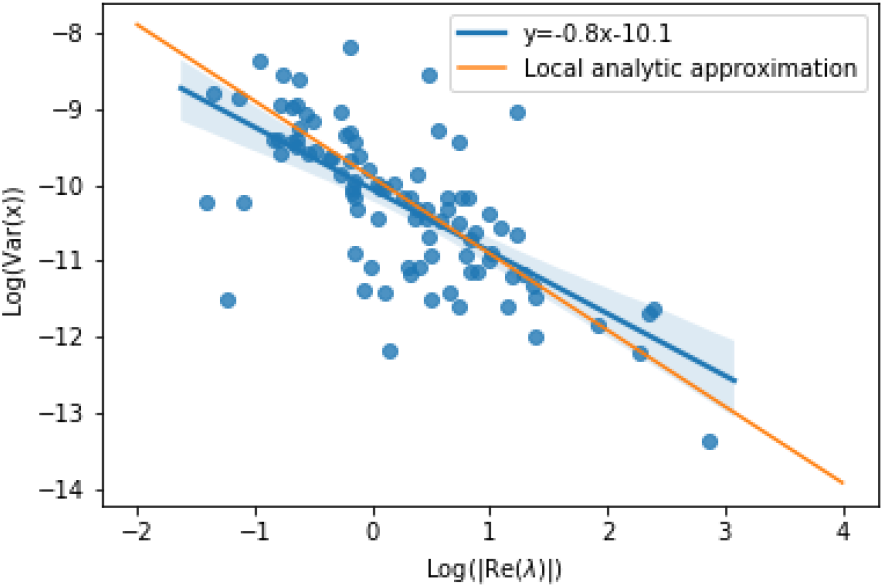
Relationship between variance of a stationary realisation and its dominant eigenvalue. The local analytic approximation is the expression in Eqn. (2). This shows good agreement with numerical simulations.

In order to control for the variance of simulations across the initial parameter configuration of each system, we need to scale *σ* accordingly. We therefore take the noise amplitude for each model to be

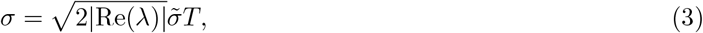

where 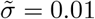 and *T* is drawn from the triangular distribution *T* ∈ T (0.75, 1, 1.25).

